# Impaired liver regeneration and lipid homeostasis in CCl_4_ treated WDR13 deficient mice

**DOI:** 10.1101/763433

**Authors:** Arun Prakash Mishra, Archana B Siva, Chandrashekaran Gurunathan, Y Komala, B Jyothi Lakshmi

## Abstract

**Background and Aim:** WDR13 - a WD repeat protein, is abundant in pancreas, liver, ovary and testis. Absence of this protein in mice has been seen to be associated with pancreatic β-cell proliferation, hyperinsulinemia and age dependent mild obesity. Previously, we have reported that the absence of WDR13 in diabetic *Lepr*^*db/db*^ mice helps in amelioration of fatty liver phenotype along with diabetes and systemic inflammation. This intrigued us to study direct liver injury and hepatic regeneration in *Wdr13*^−/0^ mice using hepatotoxin CCl_4_.

**Methods:** Mice were injected with CCl_4_ twice a week for 8 consecutive weeks. Controls were injected with vehicle (olive oil) similarly. After the last injection, mice were given a 10-days of recovery period and then sacrificed for physiological and molecular analyses.

**Results:** In the present study we report slower hepatic regeneration in *Wdr13*^−/0^ mice as compared to their wild type littermates after CCl_4_ administration. Interestingly, during the regeneration phase, hepatic hypertriglyceridemia was observed in *Wdr13*^−/0^ mice. Further analyses revealed an upregulation of PPAR pathway in the liver of CCl_4_-administered *Wdr13*^−/0^ mice, causing *de novo* lipogenesis.

**Conclusions:** The slower hepatic regeneration observed in CCl_4_ administered *Wdr13*^−/0^ mice, may be linked to liver hypertriglyceridemia because of activation of PPAR pathway.

## Introduction

Liver is a vital organ responsible for several metabolic processes and over 2 million deaths per year worldwide have been reported from liver related diseases [1]. Several factors cause liver damage, of which chronic alcohol abuse and viral hepatitis are identified as the major ones [2]. Liver, being one of the fastest regenerating organs [3], rectifies the damage and, in the repair process accumulates extracellular matrix (collagen) resulting in fibrosis [4] causing morphologically and functionally damaged liver [4]. Liver damage also occurs as a result of extensive lipid accumulation - popularly known as fatty liver, that is caused by either high fat diet intake or obesity-associated insulin resistance [non-alcohol dependent steatohepatitis or NASH] [5]. Hepatocyte damage induced by fatty liver condition leads to inflammation and fibrosis in liver [4].

To study liver damage in mouse model, chronic CCl_4_ (carbon tetrachloride) administration is an established method [6]. CCl_4_ gets metabolized to CCl_3_OO^*^ peroxide free radicals in the liver via mitochondrial cytochrome P450 (CYP450) and the generated peroxide free radicals damage the hepatocyte lipid biomembrane, through lipid peroxidation, resulting in the release of cellular contents in extracellular matrix, eliciting a myriad of inflammatory signals in liver. High level of inflammation leads to apoptosis and further liver damage [7]. These damages are, however, spontaneously reversible if the mice are given a regeneration period of 20 days [8].

WDR13, a member of the WD repeat protein family, is present in most of the mouse tissues, with relatively higher abundance in pancreas, liver, testis and ovary [9]. Previous studies have shown that *Wdr13*^*-/0*^ mice have age-dependent mild obesity, enhanced pancreatic β-cell proliferation [10–12], hyperinsulinemia and better glucose clearance [10,12]. Cellular hyperproliferation due to downregulation of p21 in absence of WDR13, has also been reported [11]. We have also demonstrated that the introgression of *Wdr13*-null mutation in obese and diabetic *Lepr*^*db/db*^ mouse model resulted in reduction of serum free-fatty acids, liver triglyceride and systemic inflammation [12]. Cell proliferative phenotype in pancreas and amelioration of fatty liver, prompted us to study the response of *Wdr13*^*-/0*^ mice to direct liver injury induced by chronic CCl_4_ administration and regeneration therein. Interestingly contrary to our previous results, we observed slower regeneration along with hypertriglyceridemia in the liver of *Wdr13*^*-/0*^ mice after CCl_4_ toxicity. The PPAR pathway was found to be upregulated in the liver of mutant mice, likely responsible for observed hypertriglyceridemia.The upregulated PPARγ could also be the plausible explanation for the slower regeneration of mutant liver. Our study thus, suggest a link between liver regeneration and lipid accumulation to be associated by PPAR pathway.

## Materials and Methods

### Animals

Institutional Animal Ethics Committee (IAEC) of CSIR-CCMB, Hyderabad, India (**Animal trial registration number**-20/1999/CPCSEA dated 10/3/99) approved the animal experiments and the experiments were performed under IAEC guidelines. 8-10 week old *Wdr13* knockout (*Wdr13*^*-/0*^) CD1 male mice [PCR genotyped as described earlier [11]] along with their wild type (*Wdr13*^*+/0*^) littermates were used in the study. Mice were housed at 22-25°C temperature with 12 hour light-dark cycle. All the mice were fed standard chow for entire experimental duration *ad libitum*.

### CCl_4_ administration

Mice were injected (intraperitoneally) with CCl_4_ (10% v/v dissolved in olive oil, 2ml/kg body weight), twice a week for 8 consecutive weeks. Controls were injected with vehicle (olive oil) similarly. After the last injection, mice were given a 10-days of recovery period and then sacrificed for physiological and molecular analyses.

### Liver/body weight ratio

Mice were weighed before sacrificing. Whole liver was carefully excised out and weighed. Ratio of respective liver weight to body weight was taken and graph was plotted.

### Histological analyses

Liver was fixed in 4% paraformaldehyde overnight and then embedded in paraffin wax. 4µm sections of livers were mounted on positively charged slides (Fischer scientific) and hematoxylin-eosin staining was performed for visualizing tissue morphology. Sirius Red staining was performed to study collagen deposition. To evaluate the number of actively dividing cells in the liver, Ki-67 immunostaining was performed according to manufacturer’s guidelines (BD Biosciences DAB substrate kit Cat.-550880, Ki-67 antibody-Millipore-AB9260). Ki-67 positive cells were counted manually (3 frames per section) from 5 different animals.

### Serum collection and analyses

500µl of blood was drawn from the mice orbital sinus and left to clot in a slanting position at room temperature for 2 hours for serum collection. The serum was collected by centrifugation at 10,000g for 10 minutes. Liver function tests-SGOT (Serum glutamic oxaloacetic transaminase) and SGPT (Serum glutamic-pyruvic transaminase) were performed as per the manufacturer’s instructions (Coral Clinical Systems). Serum glucose was tested using Accu-Chek^®^ Active Glucometer (model number: HM100005). Level of serum insulin was determined using insulin ELISA kit (Millipore-EZMRI-13K).

### TBARS assay

The assay was performed following spectrophotometeric measurement of colour generated by the reaction of MDA (Malondialdehyde) with thiobarbituric acid (TBA) as described earlier [13]. Briefly, 50mg of liver was homogenized in PBS. The supernatant was collected after centrifuging the homogenate at 10,000g. 500µl of trichloroacetic acid (10%) was added to 100µl of supernatant and heated at 95°C for 15 minutes. The mixture was cooled to room temperature and centrifuged at 3,000g for 10 minutes. 200µl of TBA (0.67% in 1M NaOH) was added to 400µl of the supernatant from the previous step. This mixture was then heated at 95°C for 15 minutes. After cooling the samples to room temperature, 200µl of the same was taken in a microtitre plate and absorbance was recorded at 532nm. MDA concentration was determined using the absorbance coefficient of the TBA-MDA complex (ε = 1.56× 10^5^ cm^−1^∙M^−1^).

### Liver triglycerides

Total lipid from liver was extracted using Folch’s method [14]. Briefly, 50mg of liver sample was homogenized in chloroform:methanol (2:1) mixture. The homogenate was vigorously agitated for 15-20 minutes and centrifuged at 12,000g. The supernatant was washed by adding 0.2 volume water and gentle mixing. The upper phase was removed and the lower one was vacuum dried. 100µl of ethanol was added to the dried tube and kept at 4°C overnight for dissolving the deposited fat. Triglyceride estimation was done using an assay kit (Coral Clinical Systems).

### Western blotting

50mg of liver sample was homogenised in lysis buffer [50mM Tris pH 8.0, 150mM NaCl, 0.1% SDS,1% Na-deoxycholate, 1% Triton X100, 1X protease inhibitor cocktail (Roche 11873580001)] and supernatant was collected after centrifuging at 10,000g. After protein estimation, 50µg of the lysate was run on SDS-PAGE and transferred to PVDF membrane. Membrane was probed with protein specific antibodies-β-actin (1:1000 dilution, Santacruz sc-47778), p53 (1:500 dilution, Santacruz sc-6243), Cyclin D1 (1:500, Santacruz sc-246), Cyclin E (1:500, Santacruz sc-481), AC1 (1:500, Novusbio NBP1-19628), p38α (1:500, Santacruz sc-535), P-p38α (1:500 dilution, Santacruz sc-101759) and PPARγ (1:500 dilution, Cell Signaling 2435S). For stripping the PVDF membrane, it was incubated in stripping buffer (0.1 M β-mercaptoethanol, 2% SDS, 62.5 mM Tris-HCl, pH 6.7) at 50°C for 30 minutes. The blot were washed thoroughly and reprobed with different primary antibodies. All the images presented here are cropped images of full length blots for better representation and all the gels were run under similar experimental conditions. ImageJ analysis of western blots was performed for quantification normalised with respective β-actin as loading control.

### RNA isolation and real-time PCR

50mg of liver sample was taken in 1ml RNAiso (Takara Cat.-9108/9109) and RNA was isolated according to the manufacturer’s guidelines. cDNA was synthesized using ImProm-II Reverse Transcription System (Promega-A3800). Quantitative real-time PCR (qPCR) was done using Ex Taq SYBR Green (Takara-RR820A), gene specific primers (Table 1) and β-actin for normalisation.

**Table 1.**
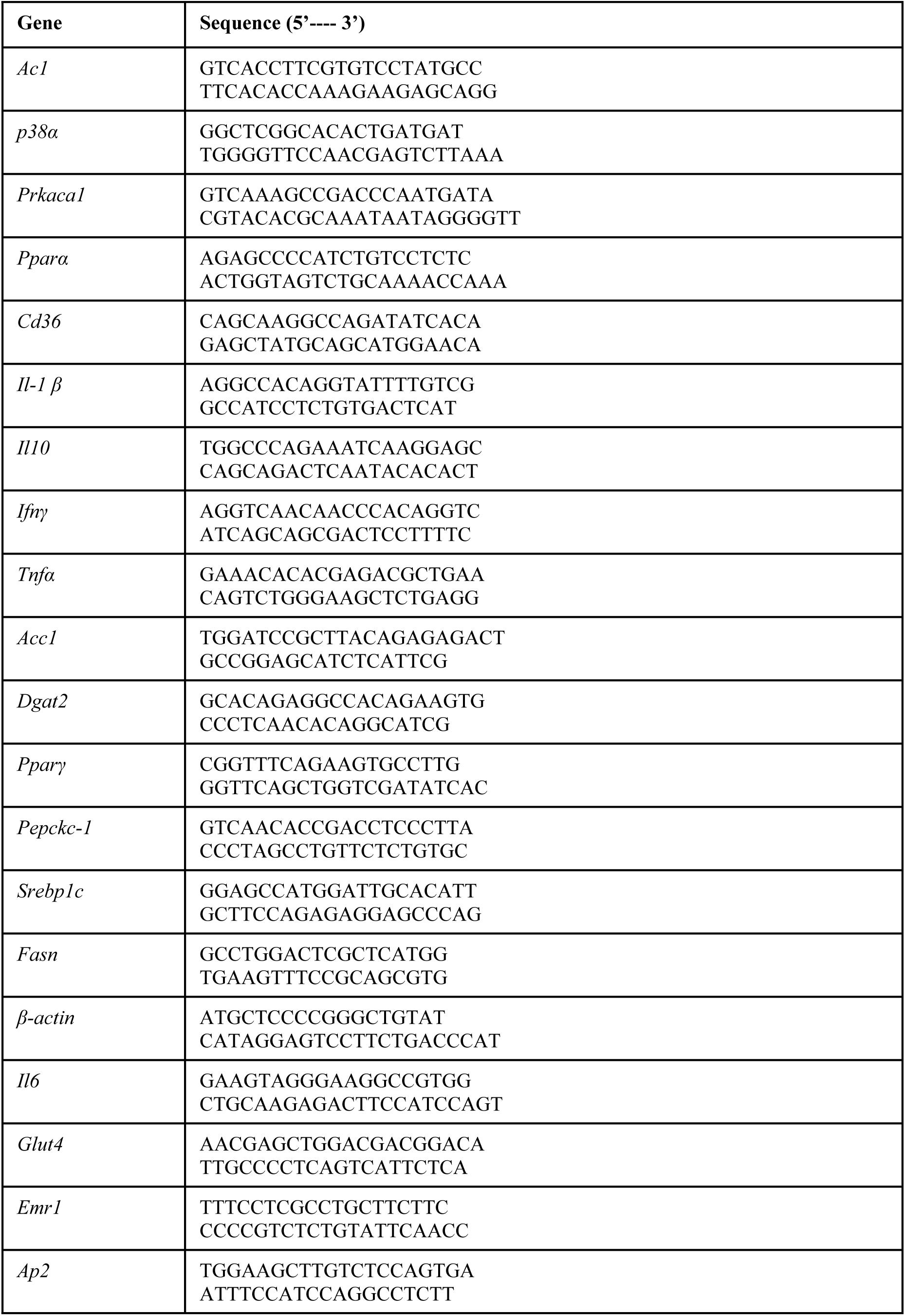
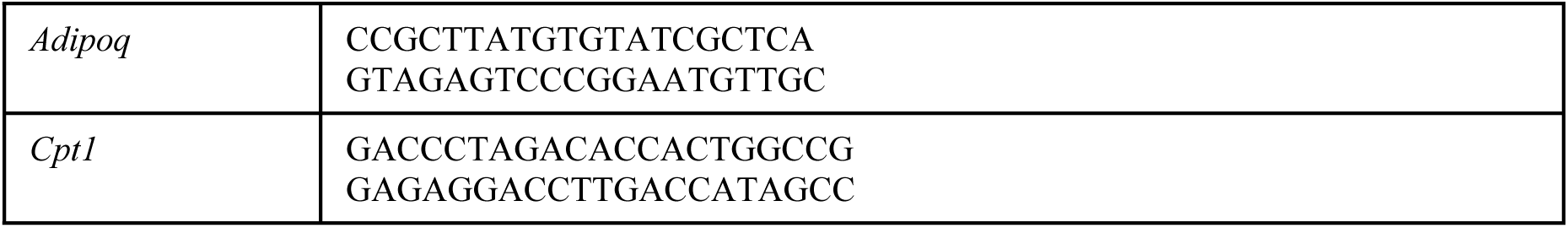
Primers used for qPCR

### Primary hepatocyte isolation and culture

Isolation of primary hepatocytes was done with the double perfusion collagenase method as described earlier [15]. Trypan blue staining was used for cell viability determination. Cells were seeded on collagen coated 6-well plates (6×10^5^ viable cells/well). The cells were grown in RPMI media (50µg/ml penicillin and streptomycin, 10% FBS) in 5% CO_2_ incubator at 37°C. After 24 hours of incubation, fresh RPMI media comprising 100µM CCl_4_ (dissolved in DMSO) was added to specific wells. After 24 hours of CCl_4_ administration, the hepatocytes were harvested and total RNA was isolated for qPCR analyses.

### CCl_4_ toxicity assay on hepatocytes

This assay was done using MTT [3-(4,5-Dimethylthiazol-2-yl)-2,5-Diphenyltetrazolium Bromide] colorimetric assay for cellular growth and survival, as described earlier [16]. Briefly, 1000 viable hepatocytes were counted and seeded in triplicates in 96-well plate (coated with collagen) and incubated in RPMI media for 4 hours for attachment. Fresh media with different concentrations of CCl_4_ (0-0.25mM dissolved in DMSO) was added to the specific wells and incubated for 24 hours at 37°C and 5% CO_2_. After 24 hours, the wells were washed with PBS and cells were incubated with 0.8 mg/ml of MTT dissolved in serum free RPMI media for 4 hours. After incubation, the wells were again washed with PBS and 200µl of DMSO was added to each well. After a gentle shaking for 10 minutes, absorbance was taken at 560nm.

### Statistical analyses

All the graphs were plotted using MS Excel worksheet. Single factor ANOVA and two-tailed unpaired student’s t-test were used for statistical analyses. Graphs represent mean ± SEM.

## Results

### Effect of *Wdr13* deletion on liver pathology and regeneration after CCl_4_ administration

CCl_4_ damages the plasma membranes of hepatocytes, which leads to increase in serum glutamic-oxaloacetic transaminase (SGOT) and serum glutamate-pyruvate transaminase (SGPT) levels [17]. The initial damage consequently elicits huge inflammatory response causing severe hepatic damage [17]. To analyse the damage caused by chronic CCl_4_ administration, we studied the liver damage parameters, namely; SGOT & SGPT, hepatocytes morphology, collagen deposition, lipid peroxidation and inflammation in WDR13 deficient mice. The mutant mice did not differ significantly from the wild type counterparts in all the above parameters (Supplementary Fig 1); though the liver/body weight ratio in *Wdr13*^*-/0*^ mice was found to be lower than that in *Wdr13*^*+/0*^ mice (Fig. 1c). The mutant mice had lower number of actively dividing hepatocytes, as revealed by Ki-67 immunostaining of the liver sections (Fig. 1a,b), indicating that the liver of *Wdr13*^*-/0*^ mice has slower regeneration. To further understand the reason for slow regeneration of *Wdr13*^*-/0*^ livers, we analysed the expression level of cell cycle genes. Lower protein levels of Cyclin D1 and Cyclin E, key molecules for cell cycle G1/S transition, and higher protein level of p53 (anti-proliferative gene) were observed in the liver of *Wdr13*^*-/0*^ mice (Fig. 1d,e).

**Figure 1.**
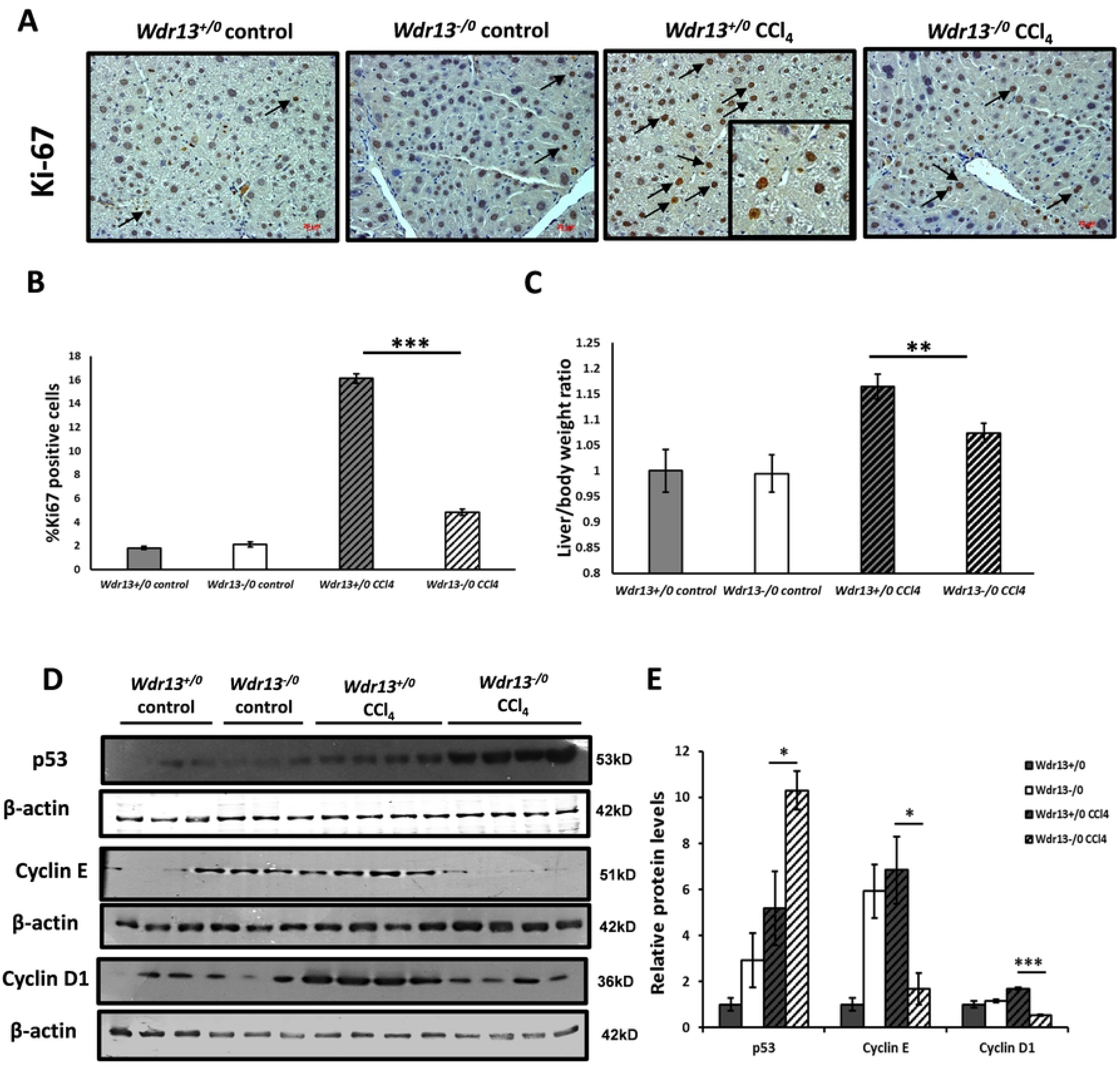
Effect of *Wdr13* deletion on liver regeneration after CCl_4_ administration. A) Ki-67 immunostaining of liver sections from control and CCl_4_ administered *Wdr13*^*+/0*^ and *Wdr13*^*-/0*^ mice. Arrows indicate positively stained nuclei (inset for better representation). B) Quantitative representation of Ki-67 positive cells from control and CCl_4_ administered *Wdr13*^*+/0*^ and *Wdr13*^*-/0*^ liver. c) Liver/body weight ratios of control and CCl_4_ administered *Wdr13*^*+/0*^ and *Wdr13*^*-/0*^ mice. D) p53, Cyclin E and Cyclin D1 immunoblots performed on liver proteins. E) Relative protein levels of p53, Cyclin E and Cyclin D1 showed in (D) immunoblots. n=5 mice for controls and n=8 mice for treatment group, **p<0.01, ***p<0.001.

### Effect of *Wdr13* deletion on liver lipid content after CCl_4_ administration

Since our previous study [12] indicated amelioration of fatty liver phenotype upon deletion of *Wdr13* in *Lepr*^*db/db*^ mice, we analysed liver triglyceride levels in the present study. In contrast to our earlier study, we observed higher triglycerides level in the liver of CCl_4_ administered *Wdr13*^*-/0*^ mice as compared to that in CCl_4_ treated wildtypes (Fig. 2b). The mutant livers were visibly pale, a characteristic feature of fatty liver, after CCl_4_ administration as compared to the wild type ones (Fig. 2A). Substantiating the above results, relative mRNA levels of lipogenic genes (*Acc1, Dgat2, Fasn*, and *Srebp1*) were also found to be upregulated in the liver of CCl_4_ administered *Wdr13*^*-/0*^ mice (Fig. 2c).

**Figure 2.**
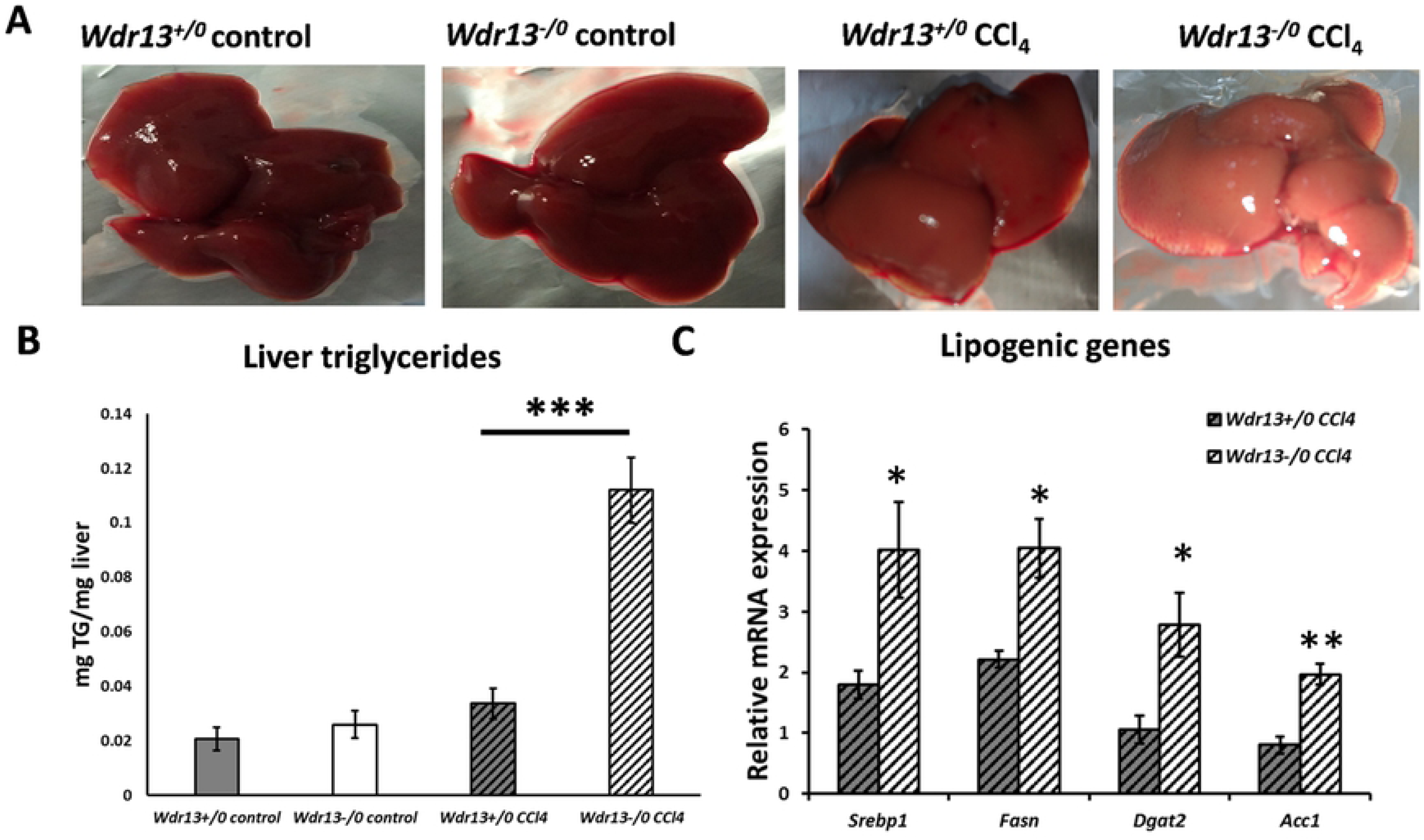
Effect of *Wdr13* deletion on liver triglyceride levels after CCl_4_ administration. A) Liver morphology in control and CCl_4_ treated *Wdr13*^*+/0*^ and *Wdr13*^*-/0*^ mice. B) Total triglyceride content of liver from control and CCl_4_ treated *Wdr13*^*+/0*^ and *Wdr13*^*-/0*^ mice. C) Relative mRNA expression levels of lipogenic genes in CCl_4_ treated *Wdr13*^*+/0*^ and *Wdr13*^*-/0*^ mice. *p<0.05, **p<0.01 and ***p<0.001.

### Analyses of lipid metabolic pathway in *Wdr13* deficient mice on CCl_4_ administration

It is known that the Peroxisome proliferator-activated receptor (PPAR) pathway plays a crucial role in lipid metabolism in liver [18] and its upregulation increases lipid accumulation therein [19–23]. We analysed the PPAR pathway proteins-PPARγ, P-p38α, p38α, and Adenylyl Cyclase 1 (AC1), by western blot analyses and observed higher levels of each of these proteins in the liver of CCl_4_ administered *Wdr13*^*-/0*^ mice (Fig. 3a, b). Additionally, the relative mRNA levels of *Pparγ* and *Pparα*, in the liver of CCl_4_ administered *Wdr13*^*-/0*^ mice, were also found to be upregulated (Fig. 3c). The downstream target genes (Fig. 3d) of PPARα & PPARγ (*Adipoq, Ap2, Cpt1* and CD36) and the upstream genes to the PPAR pathway (Fig. 3c) -*Prkaca1* (catalytic subunit of protein kinase A), *p38α* & adenylyl cyclase 1 (*Ac*1) were all upregulated, confirming higher activity of PPARs in the liver of CCl_4_ administered *Wdr13*^*-/0*^ mice via the p38α/MAPK14 and PKA pathway. This upregulation of PPAR pathway in CCl_4_ treated *Wdr13*^*-/0*^ livers leads to the *de novo* lipogenesis and accumulation of triglycerides therein.

**Figure 3.**
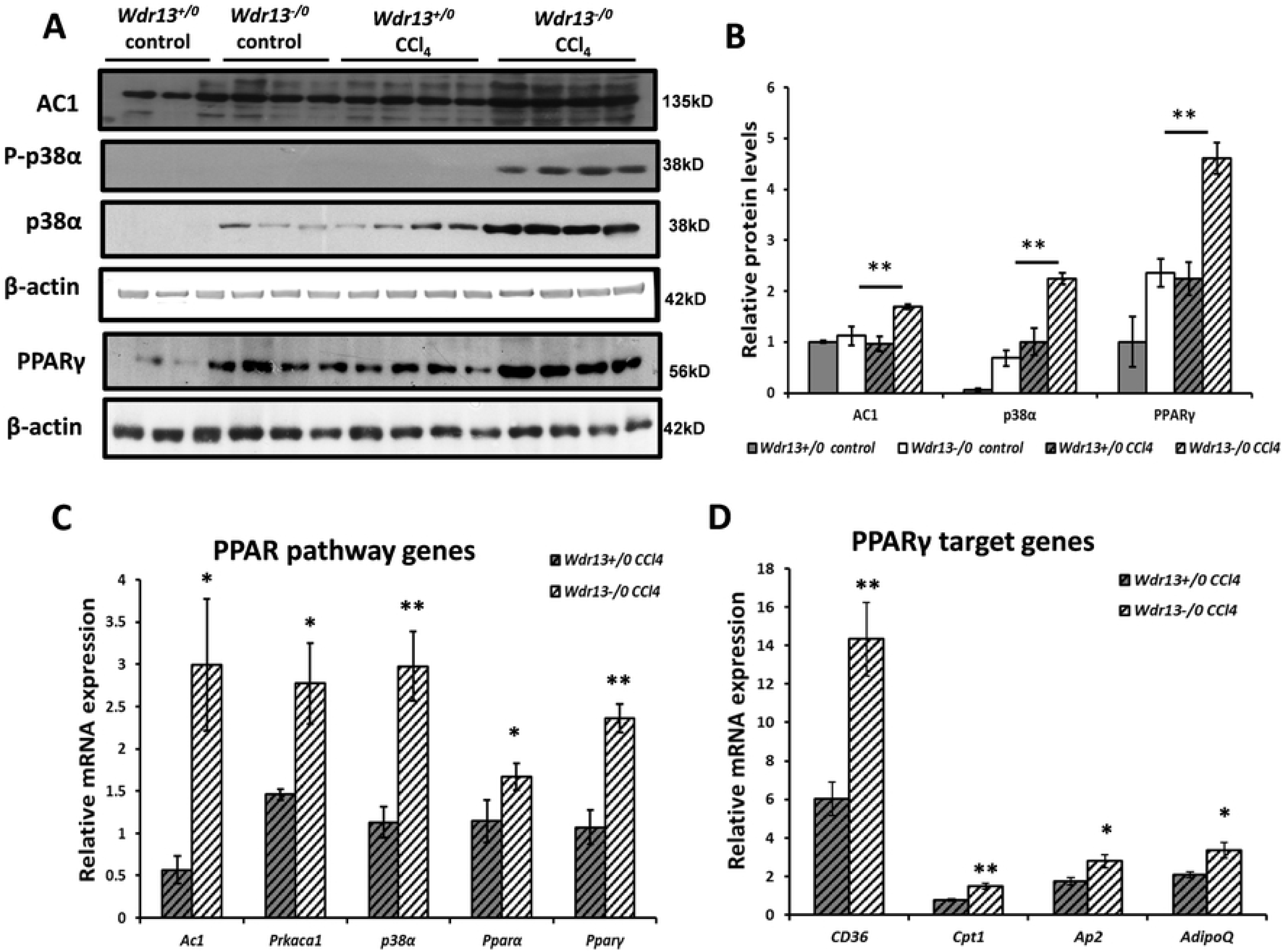
Effect of *Wdr13* deletion on liver PPAR pathway after CCl_4_ administration. A) AC1, P-p38α, p38α and PPARγ immunoblots performed on liver proteins. B) Relative protein levels of AC1, p38α and PPARγ showed in (A) immunoblots. C) Relative mRNA expression levels of PPAR pathway genes in CCl_4_ treated *Wdr13*^*+/0*^ and *Wdr13*^*-/0*^ mice. D) Relative mRNA expression levels of PPARγ target genes in CCl_4_ treated *Wdr13*^*+/0*^ and *Wdr13*^*-/0*^ mice. n=5 mice for controls and n=8 mice for treatment group, *p<0.05, **p<0.01. (AC1-Adenylyl Cyclase 1)

### CCl_4_ toxicity on primary hepatocytes

The mouse model under the present study is a whole body *Wdr13* gene knockout, and therefore, the observed phenotype may have resulted because of the systemic absence of WDR13. To understand if the observed phenotype was indeed due to the absence of WDR13 in liver, we performed experiments with primary hepatocytes from *Wdr13*^*+/0*^ and *Wdr13*^*-/0*^ mice. CCl_4_ toxicity on primary hepatocytes was analysed by culturing the isolated hepatocytes with increasing concentration of CCl_4_ (0.0 - 0.25mM). MTT assay revealed that *Wdr13*^*-/0*^ hepatocytes are more susceptible to CCl_4_ (Fig. 4a), as seen by reduced cell viability at even the lowest concentration of CCl_4_ (0.05mM).

**Figure 4.**
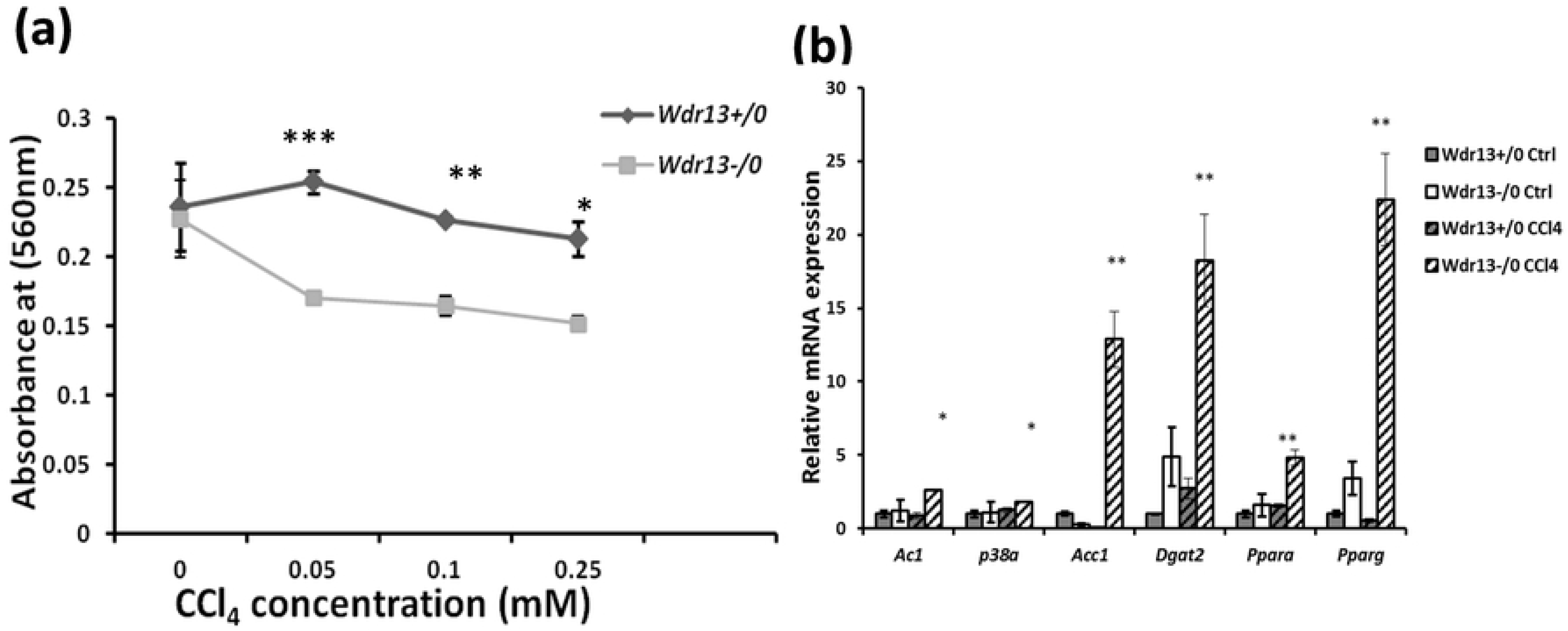
Effect of *Wdr13* deletion on CCl_4_ toxicity in primary hepatocytes. A) CCl_4_ toxicity on primary hepatocytes analysed by MTT assay. B) Relative mRNA expression levels of genes involved in lipid metabolism studied in hepatocytes treated with 100µM CCl_4_ for 24 hours. *p<0.05, **p<0.01, ***p<0.001, n=3.

To analyse the genes in the PPAR pathway in primary hepatocytes, *Wdr13*^*+/0*^ and *Wdr13*^*-/0*^ hepatocytes were treated with 0.1M CCl_4_ for 24 hours and then harvested for total RNA isolation and qPCR. Consistent with the *in vivo* results, the qPCR analysis of cDNA from CCl_4_ treated hepatocytes indicated upregulation of the PPAR pathway along with the upstream genes-*Ac1* & *p38α* and other lipid metabolism genes (*Acc1* & *Dgat2*) in *Wdr13*^*-/0*^ hepatocytes (Fig. 4b).

### Effect of *Wdr13* deletion on serum parameters and levels of insulin responsive genes in liver after CCl_4_ administration

Hyperglycemia and hyperinsulinemia are one of the major causes for fatty liver [24]. We analysed random serum insulin and glucose levels in control and CCl_4_ administered *Wdr13*^*+/0*^ and *Wdr13*^*-/0*^ mice and observed no significant difference in these parameters (Fig. 5a,b). Obesity-induced insulin resistance is considered to be a prime inducer of hepatosteatosis [25]. Thus, we examined insulin responsive genes in the liver, namely, *Glut4* and *Pepckc-1* by qPCR and their levels were found to be similar in CCl_4_ administered *Wdr13*^*+/0*^ and *Wdr13*^*-/0*^ mice (Fig. 5d). *Cpt1*, a key gene participating in β-oxidation of lipids [5] that gets inhibited by hyperglycaemia and hyperinsulinemia [25] was upregulated in *Wdr13*^*-/0*^ livers (Fig. 5d). Lipolysis and dyslipidemia are yet major factors leading to fatty liver [5]. We thus analysed serum triglycerides and cholesterol levels in control and CCl_4_ administered the *Wdr13*^*+/0*^ and *Wdr13*^*-/0*^ mice and found no significant differences (Fig. 5c). Taken together, these results indicated that the observed hepatic hypertriglyceridemia was more likely due to *de novo* lipogenesis in *Wdr13*^*-/0*^ mice rather than due to systemic absence of *Wdr13* gene in tissues other than the liver.

**Figure 5.**
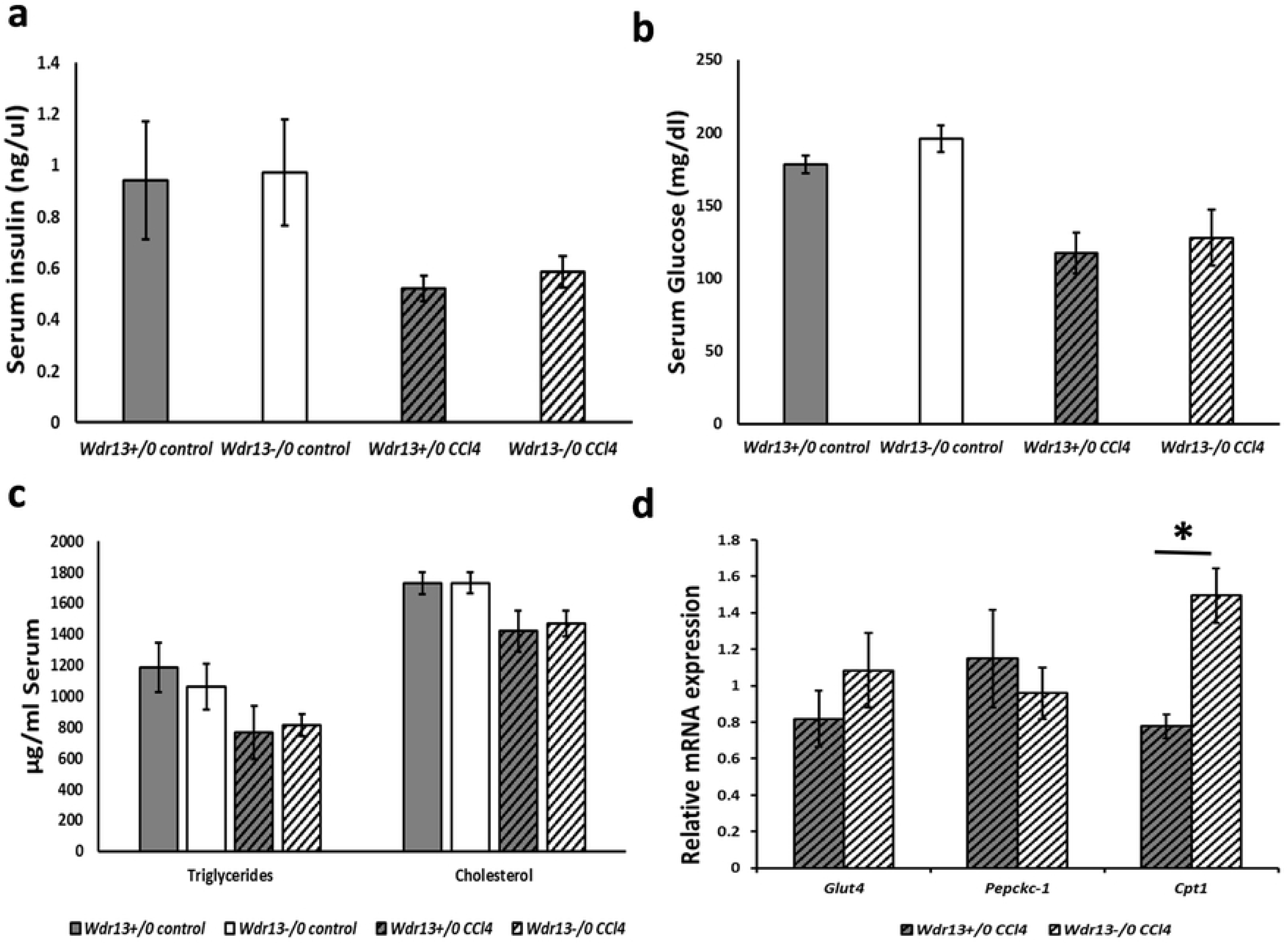
Effect of *Wdr13* deletion on serum insulin, glucose, lipids levels and mRNA expression levels of insulin responsive genes in liver after CCl_4_ administration. A) Mean random serum insulin levels (ng/µl) in control and CCl_4_ administered *Wdr13*^*+/0*^ and *Wdr13*^*-/0*^ mice. B) Mean random serum glucose levels (mg/dl) in control and CCl_4_ administered *Wdr13*^*+/0*^ and *Wdr13*^*-/0*^ mice. C) Serum triglycerides and cholesterol levels (µg/ml) in control and CCl_4_ administered *Wdr13*^*+/0*^ and *Wdr13*^*-/0*^ mice. D) Relative mRNA expression levels of insulin responsive genes in CCl_4_ administered *Wdr13*^*+/0*^ and *Wdr13*^*-/0*^ liver. n=5 mice for controls and n=8 mice for treatment group, *p<0.01

## Discussion

CCl_4_ is an established hepatotoxin to study liver damage and regeneration [6]. It damages the liver initially via its conversion to CCl_3_OO^*^ free radical by Cytochrome P450 and later by eliciting myriad of inflammatory signals [7]. In the present study, liver damage was induced in *Wdr13*^*+/0*^ and *Wdr13*^*-/0*^ mice by challenging them with CCl_4_. The *Wdr13*^*-/0*^ mice exhibited lower number of regenerating hepatocytes as compared to *Wdr13*^*+/0*^ mice, thus leading to lower liver/body weight ratio. Also CCl_4_ was found to be more toxic on *Wdr13*^*-/0*^ primary hepatocytes as compared to *Wdr13*^*+/0*^ hepatocytes.

*Wdr13*^*-/0*^ livers have higher p53 and lower Cyclin D1 & E expression after CCl_4_ administration, plausibly attributed to the elevated PPARγ levels. It is known that PPARγ activation slows the liver regeneration process [26], upregulates p53 levels (anti-proliferative gene) and downregulates cyclins in the liver [27,28]. This seems to be an obvious reason for slower liver regeneration and lower liver/body weight ratio in *Wdr13*^*-/0*^ mice.

We also observed that while regenerating from chronic CCl_4_ toxicity, *Wdr13*^*-/0*^ mice accumulated higher amounts of triglycerides in liver, a condition seen in fatty liver or hepatosteatosis, seen to be via the PPAR (PPARα and PPARγ) pathway. The role of PPARs in adipose lipid metabolism and liver lipid homeostasis is well established [18,22]. Literature reveals that liver-specific overexpression of *Pparγ* induces lipogenic gene expression, leading to hepatosteatosis [19]. Similarly, liver-specific deletion of *Pparγ* protects mice from hepatic lipid accumulation [20]. Studies also show PPARα to be a key protein involved in liver lipid metabolism [22] and its upregulation results in *de novo* lipogenesis and lipid chain elongation in liver [29]. The present study emphasises role of PPAR (PPARα and PPARγ) pathway in CCl_4_ induced hepatosteatosis in *Wdr13*^*-/0*^ mice.

p38α/MAPK14 plays a pivotal role in PPAR signalling [30,31] as it upregulates the transcriptional activity of *Pparγ* in mouse adipocytes [32] and human trophoblasts [33]. PPARs are directly linked to cAMP signalling pathway via p38α/MAPK14 [23,31,32,34]. In the present study, the activation of p38α/MAPK14 along with its regulatory genes-protein kinase A (PKA) and adenylyl cyclase 1 (Fig. 3), in liver of CCl_4_ treated *Wdr13*^*-/0*^ mice, determines its role in PPAR (PPARα and PPARγ) pathway upregulation. This indicates that under CCl_4_ stress and WDR13 absence, PPAR pathway gets activated. Our previous study [12] has also indicated the role of WDR13 in regulation of *Pparγ* expression. Besides hepatotoxin stress, this study also reveals that the absence of WDR13 *per se* renders the liver susceptible to steatosis owing to the upregulated PPARγ and p38α in the control (vehicle treated) *Wdr13*^*-/0*^ mice (Fig. 3a,b). However, further investigations are required to delineate the exact role of WDR13 in the regulation of the said pathway.

Previously, we studied the liver physiology in *Lepr*^*db/db*^ and *Wdr13^−/0^*|*Lepr*^*db/db*^ double knockout mice [12]. Onset of fatty liver in *Lepr*^*db/db*^ mice is due to lipolysis and dyslipidemia resulting in heavy load of triglycerides and free-fatty acids in serum [35]. WDR13 deletion in *Lepr*^*db/db*^ mice improved adipose and pancreatic function which resulted in reduced dyslipidemia and hyperglycemia, two systemic factors contributing towards hepatosteatosis, leading to reduced serum triglycerol and free-fatty acids hence ameliorating the fatty liver phenotype. We did not study the expression of *Pparγ* in liver in the previous study, but other lipogenic genes were found to be upregulated in *Lepr*^*db/db*^ mice in comparison to that in double knockout mice. The study confirmed that the betterment of hepatic lipid profile in double knockout mice was due to improvement in the systemic factors as discussed above. Unlike the results of previous study, here we observed hepatic hypertriglyceridemia in *Wdr13*^*-/0*^ mice during regeneration after CCl_4_ intoxication. This hepatic hypertriglyceridemia is likely due to *de novo* lipogenesis via activated PPAR pathway. Activation of PPAR pathway is also observed in *Wdr13*^*-/0*^ primary hepatocytes when treated with CCl_4_, which confirms that hypertriglyceridaemia seen *in vivo* is indeed due to the liver-specific absence of WDR13. The circulating triglycerides and cholesterol levels in *Wdr13*^*-/0*^ mice were also found similar to that in *Wdr13*^*+/0*^ mice, suggesting that the observed liver hypertriglyceridemia is indeed due to *de novo* lipogenesis and not because of systemic factors.

Hepatosteatosis is often associated with hyperglycemia and hyperinsulinemia [24]. However, in our study, there was no difference in serum insulin and glucose levels in control and CCl_4_ administered *Wdr13*^*+/0*^ and *Wdr13*^*-/0*^ mice. Consistent with these data, the relative mRNA expression levels of insulin responsive genes (*Glut4* and *Pepck-c*) were also similar in the liver of CCl_4_ administered *Wdr13*^*+/0*^ and *Wdr13*^*-/0*^ mice. Another insulin responsive gene, *Cpt1* which is required for mitochondrial β-oxidation of lipids [5] was upregulated in the liver of mutant mice. It may be noted that the expression and activity of *Cpt1* is PPARα dependent [22] and the increased expression of hepatic *Cpt1* is seen in fatty liver and obesity [36]. The upregulation of *Cpt1* in *Wdr13*^*-/0*^ liver may be a result of upregulated *Pparα* and liver hypertriglyceridemia.

Many genome wide studies show dysregulation of WDR13 in hepatic disorders. WDR13 was found to be down regulated and aided in lipogenesis in human hepatocytes [37]. Also the expression levels of WDR13 was found to be altered in hepatic oxidative stress and steatosis conditions via BIM activation [38]. Other studies too indicate indirect dysregulation of WDR13 in *in-silico* or obesity conditions [39],[40]. All these studies along with the present study reveal that loss of WDR13 per se increases the PPARγ expression.

Our recent data suggests that WDR13 functions via a multi protein complex c-Jun/NCoR1/HDAC3 and it acts as a transcriptional activator of AP1 target genes [41]. Besides, our previous studies have shown that the lack of WDR13 enhances the cell cycle and overexpression of the same inhibits cell proliferation [11]. It is possible that WDR13 may regulate Cyclin E expression via AP-1 and c-Jun/NCoR1/HDAC3; although presently we do not have direct evidence of it. It may be emphasised that the *Wdr13*^*-/0*^ mice display hyperinsulinemia and mild obesity around 12 months of age without any evidence of fatty liver phenotype [11]. To avoid confounding effects of the absence of WDR13 *per se* on metabolism and the toxic effects of CCl_4_, the present study was conducted at the age of 8-10 weeks old animals when there is no onset of hyperinsulinemia and obesity. Fatty liver condition in mice has been reported upon treatment with CCl_4_ along with high fat diet [42] via upregulation of PPARγ in the liver [43]. In this study we observed fatty liver phenotype in *Wdr13*^*-/0*^ mice on CCl_4_ treatment alone, owing to upregulation of PPAR pathway. Thus in our opinion, this study introduces WDR13 as a molecular factor, the absence of which predisposes mice to hepatosteatosis in the presence of CCl_4_ (without the requirement of high fat diet).

Summarily, in the present study we report that the *Wdr13*^*-/0*^ mice are more susceptible to CCl_4_-induced hepatotoxicity as compared to their wild type counterparts. Also the *Wdr13*^*-/0*^ mice exhibit liver hypertriglyceridemia due to *de novo* lipogenesis, during the regeneration phase after CCl_4_ toxicity. PPAR pathway upregulation in liver has a key role in the observed phenotypes of CCl_4_ administered *Wdr13*^*-/0*^ mice. CCl_4_ is an established hepatotoxin to study hepatic damage, fibrosis and regeneration in murine models, but fatty liver has never been reported in exclusive CCl_4_ toxicity in mice. *Wdr13*^*+/0*^ mice manage the CCl_4_ challenge and do not allow PPAR pathway to get upregulated. This study indicates that under CCl_4_ stress PPARγ gets overexpressed in the absence of WDR13 suggesting WDR13 may have a role in regulation of PPARγ expression, as also seen in our previous study [12]. We want to highlight the fact that absence of WDR13 *per se* does not induce *de novo* lipogenesis and fatty liver but when these mutant mice are subjected to CCl_4_ stress, liver hypertriglyceridemia is observed via upregulation of PPAR pathway. However, at present we do not understand how the absence of WDR13 activates the PPAR pathway. Further experiments are required to elaborate the mode of action of WDR13 in regulation of PPARs.

## Acknowledgements

APM, CG and ABS conceived the idea, designed experiments and wrote the manuscript. APM, YK and BJL performed the experiments. We thank Mr. Avinash T Raj for histology and Dr. Sesikeran for histological assessments. We acknowledge the Council of Scientific and Industrial Research (CSIR), New Delhi, India for financial support to Mr Arun Prakash Mishra. The experiments were conducted in Dr Satish Kumar’s laboratory.

No separate funding was allotted for this study. Authors declare no conflict of interest.

